# Heading direction with respect to a reference point modulates place-cell activity

**DOI:** 10.1101/433516

**Authors:** P.E. Jercog, Y. Ahmadian, C. Woodruff, R. Deb-Sen, L.F. Abbott, E.R. Kandel

## Abstract

Utilizing electrophysiological recordings from CA1 pyramidal cells in freely moving mice, we find that a majority of neural responses are modulated by the heading-direction of the animal relative to a point within or outside their enclosure that we call a reference point. Our findings identify a novel representation in the neuronal responses in the dorsal hippocampus.

## Introduction

Successful navigation requires knowledge of both location and direction, quantities that must be computed from self-motion and environmental cues. Place cells in the dorsal hippocampus (O’Keefe and Dostrovsky 1971), grid cells in entorhinal cortex (Fyhn et al 2004), and head-direction cells in various brain regions (Wiener et al 1989, Taube et al 1990, Mueller et al 1994, Sargolini et al 2006, Rubin et al 2014) are leading candidates for providing neural representations of location and direction. Recent studies have revealed goal-oriented cells hippocampus of the bat tuned to the direction of motion relative to rewarded locations within an environment (Sarel et al 2017; see also Hoydal et al 2018). Here, we provide evidence that CA1 neurons in mice exhibiting selectivity for location are also modulated by heading direction relative to a reference point located inside or outside their enclosure.

## Results

We recorded 1244 neurons in the dorsal region of both hippocampi (area CA1;Methods) in 12 C57BL/J6 mice recorded daily during 4 sessions of 10 minutes each over a period of 10 days. The analysis presented here is based on 697 neurons that satisfied a set of minimal response and sampling criteria (Methods). During each session, food deprived animals foraged freely within a 50 by 50 cm enclosure. In addition to monitoring the location of the animal (x, y), we tracked its absolute heading-direction angle (H), defined as heading direction relative to a fixed orientation in the room (coinciding with a cue card on one wall). The enclosure was divided into 100 square bins (5 cm by 5 cm each) and the heading angle into 10 bins of 36° each. Firing rates of the recorded neurons were computed for each bin, yielding firing-rate data r(x, y, H).

**Figure 1:**
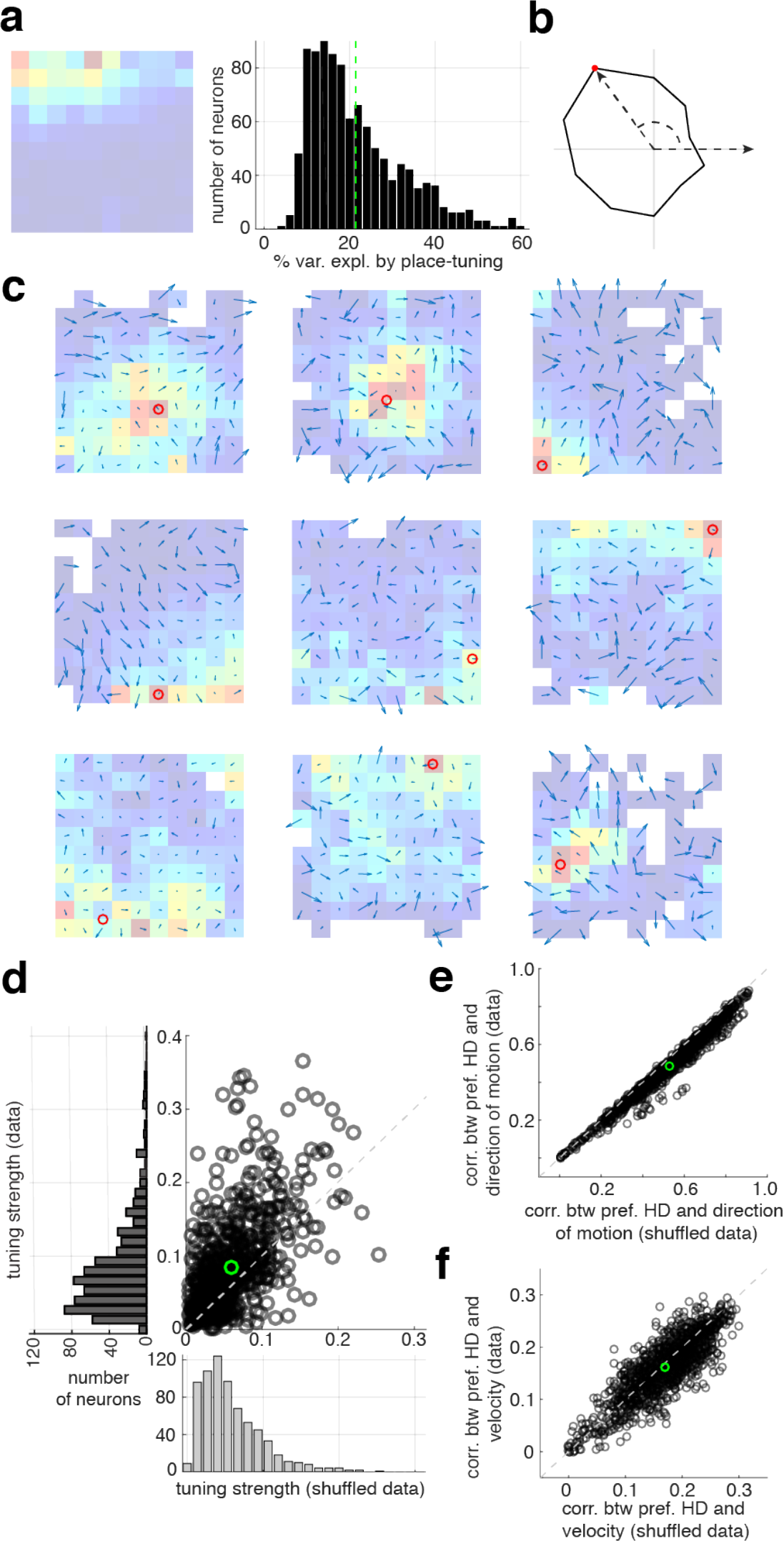
Heading-direction modulation. **a)** Firing rate as a function of location for a typical place cell (left). Colors from blue to red indicate low to high firing rates. Percent variance across neurons explained by location tuning (right). **b)** Example of heading-direction tuning in one neuron. Radius of curve is proportional to the firing rate. **c)** Tuning of firing rate for heading direction shown as rate-weighted sums of heading direction vectors within each bin. Place fields are also shown, as in a. Red circles are the centers of mass of the place fields. White squares denote bins with average firing rates too low to be included in the analysis of directional modulation. **d)** Average tuning strengths of heading-direction modulation across bins for actual and randomly reshuffled data. Tuning strength is defined as the lengths of vectors computed as in c, averaged over bins for each neuron. For surrogate data, head-direction angles were randomly reshuffled in each bin. Green circle denotes the means of both distributions. **e, f)** Correlation between the preferred heading direction (pref. HD) and either the direction of motion (top) or the velocity (bottom) for actual compared to shuffled data.

Responses of most of the recorded neurons were modulated by the location of the animal, and many would be classified as place-cells (Fig.1a (left)). Additional modulation of the activity of hippocampal neurons by heading direction relative to a fix orientation within the experimental room has been reported (Fig.1b Wiener et al 1989, Rubin et al 2014), although confounds associated with animal behavior have challenged the interpretation of this effect (Mueller et al 1994). Because place tuning accounts for a relatively small percentage of the variance of our recorded neurons (less than 50%, and often considerably less;Fig.1a (right)), we investigated modulation of the neuronal responses by heading direction defined more generally, that is, relative to various points in the environment.

A conventional place-cell description is obtained by averaging the rates r(x, y, H) over heading angles yielding a place-modulated rate r(x, y). To examine directional modulation, we considered the ratio R(x, y, H) = r(x, y, H)/r(x, y) for all bins that have firing rates greater than 0.5 Hz (we eliminated smaller rates to avoid numerical problems with small denominators; results did not change when we changed this cutoff). We multiplied unit vectors corresponding to angles centered within each heading-direction bin by the associated rate ratio R(x, y, H) and averaged to compute a preferred heading vector for each bin (<u>Fig.1c</u>; <u>Methods</u>; for further examples see <u>SupplFig.1</u>). We quantified heading-direction tuning by computing the lengths of these preferred heading vectors and averaging them over bins to obtain a tuning strength for each neuron. We then compared tuning strengths computed from the data with results obtained by randomly shuffling the heading angles within each bin (Methods). In this comparison, 70% of the recorded neurons have stronger modulation by heading direction for the data than for the shuffled data, a highly statistically significant effect (Fig.1d; p<<0.001).

We next tested if the heading-direction dependence arose from a lack of uniformity in angular sampling due to the behavior of the mice. We found, to the contrary, that omitting from the analysis spatial bins with poor angular sampling (visited with a biased distribution of angular trajectories) actually increases heading-direction tuning (SupplFig.2). In addition, we computed correlations of the average direction of motion or velocity for each bin and neuron with the preferred heading direction vectors as defined in figure 1b (Fig.1e,f). These correlations show a small but significant (p<<0.001, p < 0.03, respectively) decrease for real compared to shuffled data, which is the opposite direction from what is needed to account for the neural tuning. Finally, the middle row of figure1c shows 3 neurons that were recorded simultaneously and thus have identical behavioral effects. Nevertheless, these 3 neurons have different preferred directional tunings.

Having demonstrating heading-direction tuning per spatial bin, we went on to characterize the structure of this tuning across the spatial extent of the enclosure. To construct a more general model than simple modulation by heading relative to a fixed direction in the room, and inspired by recent work (Sarel et al 2017), we considered firing-rate modulation as a function of the angle (*θ*) between the heading direction and a line extending from the animal to a fixed but, initially, arbitrary location in the room, either within or outside the enclosure (Fig.2a). We call this fixed location the reference point and the angle thus defined the reference-heading (RH) angle. Specifically, we considered the model R(x, y, H) = 1 + g*cos(*θ* - *θ*_p_) with 4 fitting parameters for each neuron: g, the modulation strength, *θ*_p_, the preferred reference-heading angle, and two parameters describing the location of the reference point. The angle *θ* depends on x, y, H as well as on the location of the reference point. We used a subtraction procedure to assure that the average of the modulation over H is equal to 1 to remove biases (Methods).

Tuning strengths, defined as in figure 1d, were computed by multiplying the rates r(x, y) by the modeled direction modulation, and these were then compared between actual and shuffled data. A majority of neurons (70%) showed stronger tuning for actual than shuffled data (Fig.2b). We also computed the percent variance accounted for by the modulated rates and compared it to the percentages accounted by the direction-unmodulated rates r(x, y). Many neurons showed a substantial increase in explained variance when the directional-modulation described by the model was included (Fig.2c&d). This model describes the directional tuning of many neurons quite accurately (Fig.2e).

**Figure 2:**
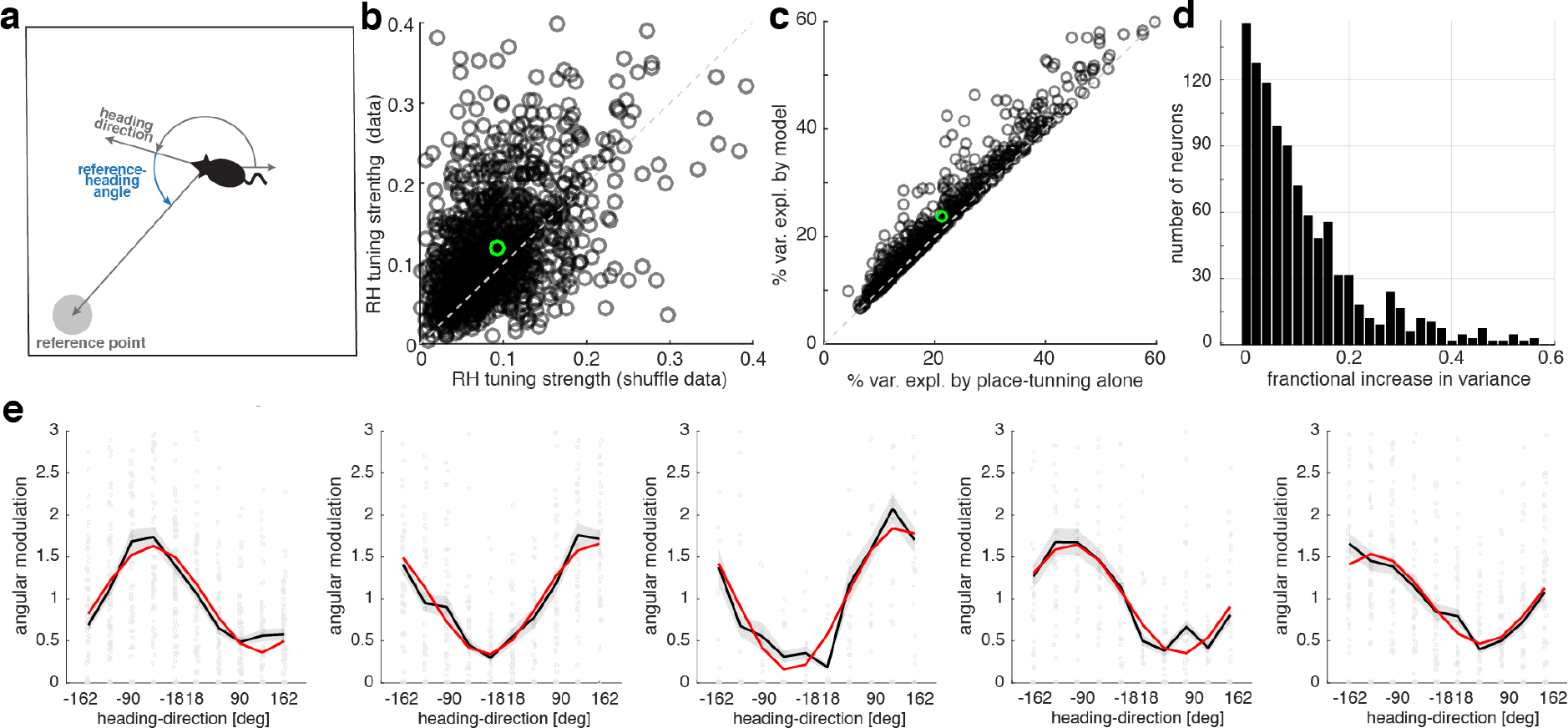
Model of modulation by heading-direction relative to a reference point. **a)** Definitions of the reference point and reference-heading angle. **b)** Reference-heading (RH) angle tuning for the recorded population of neurons compared to the tuning for surrogate data with randomly shuffled heading directions. Green circle is defined as in figure 1. **c)** Variance explained by the model with both reference-heading angle and place modulation compared to a purely spatial description. **d)** Histogram of the fractional change in variance, defined as the change in variance divided by the variance explained solely by place modulation. **e)** Examples of model fit (red line) across heading-angle bins for 5 different neurons. Black lines are means and shaded areas are standard error of the mean for the data. The red line is an average of the model over spatial bins. Grey points show all the data for each neuron.

The fitting process provides a distribution of the modulation strengths (g; Fig.3a), the preferred reference-heading directions (*θ*_p_; Fig.3b), and the x and y coordinates of the reference point (Fig.3c). The firing rates of most of the neurons are maximally enhanced when the animal moves toward the reference point (*θ*_p_ = 0). Reference points for the majority of the neurons (60%) are inside the enclosure, 8% are outside but close to the enclosure, and 32% are distal (Fig.3d). Neurons with distal reference points are effectively modulated by the absolute heading direction of the animal (Suppl.Fig.3). For neurons with reference points inside the enclosure, we found no correlation between the location of the reference point and the center of the place field (Fig.3e).

We also utilized an independent method to search for reference point locations based on an iterative algorithm that finds the centers of divergences and curls from the heading direction responses vector fields (Suppl.Fig.4). We found that the points of maximum divergence and curl generated by this method provide a good estimation of the reference point locations fitted by the model, coinciding in a majority of the cells within the same quadrant of the enclosure (Suppl.Fig.5).

**Figure 3:**
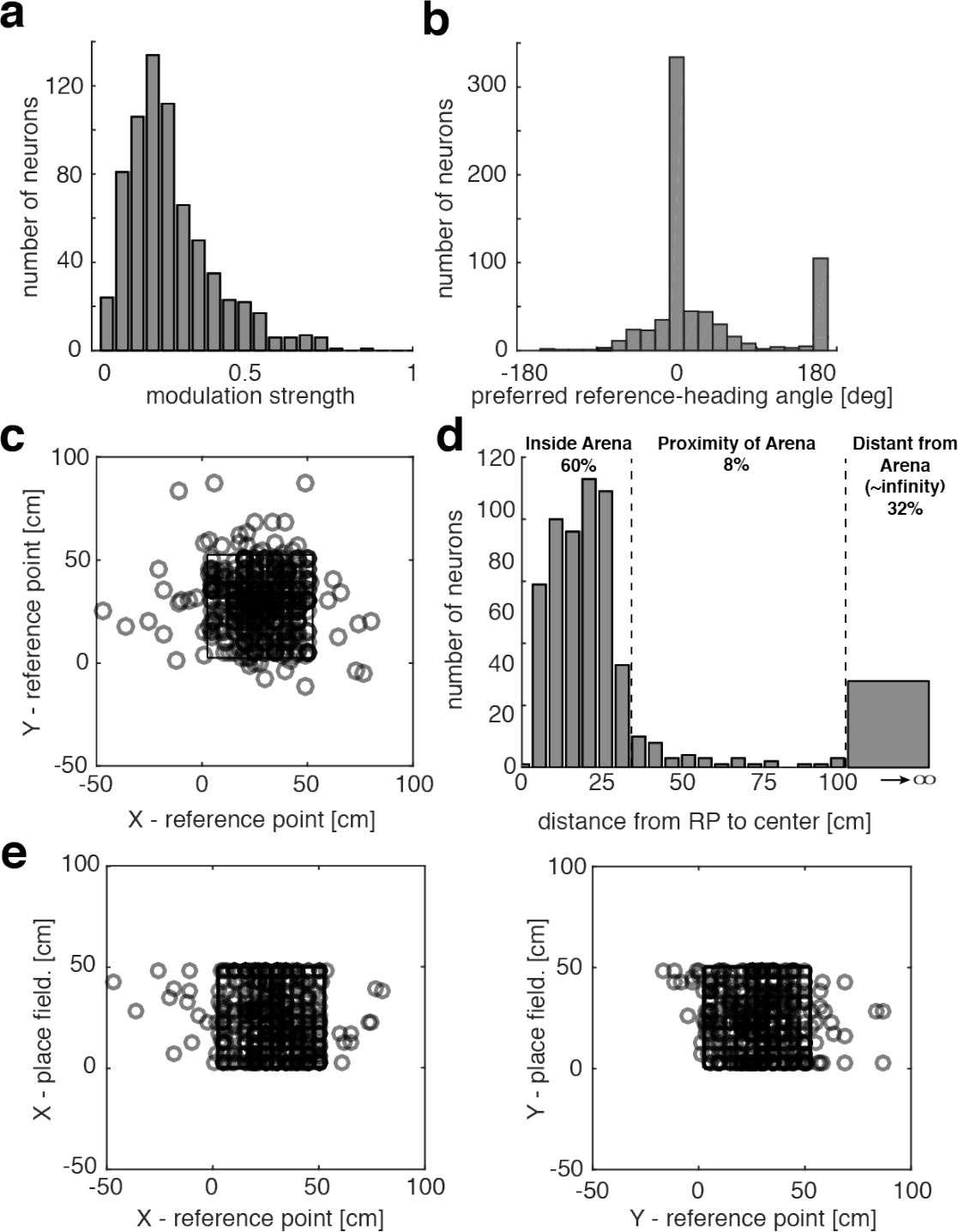
Fitted model parameters across neurons. **a)** Histogram of the amplitude of the angular modulation. **b)** Histogram of angle of the preferred reference-heading direction. **c)** Reference point locations inside and outside the enclosure. The the square in the center of the panel indicates the enclosure. Distal reference points do not appear in this figure. **d)** Histogram of the distances of reference points (RP) from the center of the enclosure. **e)** Coordinates of the reference points (x-left, y-right) compared to the coordinates of the centers of the place fields for each neuron.

## Conclusions

We find that the firing rates of the majority of neurons in the CA1 region of the dorsal hippocampus are modulated by the heading direction of the animal relative to a reference point in the environment. The majority of the neurons have a higher response when the animal moves towards the location of the reference point. This generalizes the idea of goal-(Sarel et al 2017) and object-oriented (Hoydal et al 2018) neurons to a more abstract relationship of heading direction to an unmarked point in the environment, the reference point, and it accounts for previously reported head-direction cells (Wiener et al 1989) as neurons with reference points located far from the enclosure.

## Acknowledgments

PEJ’s research supported by the Mathers Foundation and Swartz Program in Computational Neuroscience.

YA’s research supported by the Swartz Program in Computational Neuroscience.

LFA’s research supported by NSF NeuroNex Award DBI-1707398, the Gatsby Charitable Foundation and the Simons Collaboration for the Global Brain.

ERK’s research supported by the Mathers and Howard Hughes Medical Institute.

## Contributions

PEJ designed the experiment, did the surgeries for the recordings, analyzed the data and helped write the manuscript; YA helped designing statistical tests; CW ran behavioral experiments; RD performed data pre-processing; LFA developed the model, analyzed the data and wrote the manuscript; ERK helped with the design of the experiment and writing of the manuscript and supervised the project.

## Methods

### Subjects

Data was recorded from 12 male C57BL/6J mice. In 4 mice, 1 microdrive with a bundle of 4 tetrodes was implanted in the right hippocampal area CA1, and the other 8 mice were implanted with two microdrives each, with 4 tetrodes in both hemispheres (area CA1: 1.8mmDL, 1.8mmRC, ~1.1mm deep). Mice were 6-9 month old (30-34g.) at the time of the implantation surgery. After surgery, all mice were individually housed under 12 hr light/dark cycle and provided water *ad libitum* and one food pellet overnight (maintaining the same body-weight as prior to the implantation (+ the implant)). All housing procedures conformed to National Institute of Health (NIH) standards using protocols approved by the Institutional Animal Care and Use Committee (IACUC).

### Electrode implantation and surgery

Custom-made, reusable microdrives (Axona) were constructed by attaching an inner (23 ga) stainless steel cannula to the microdrive’s movable part to allow inserting the electrode reliably at 10 microns steps. An outer (19 ga) cannula covered the exposed tetrode section from the dura to the inner cannula. Tetrodes were built by twisting four 17 μm thick platinum-iridium wires (California wires) and then heat bonded them. Four such tetrodes were inserted into the inner cannula of the microdrive and connected to the wires of the microdrive. Prior to surgery, the tetrodes were cut to an appropriate length and plated with a platinum/gold solution with additives (Ferguson J.E. et al 2009) until the impedance dropped to 80-100 KΩ. All surgical procedures were performed following NIH guidelines in accordance with IACUC protocols. Mice were anesthetized with a mixture of 0.11 ml of Ketamine and Xylazine (100 mg/ml, 15 mg/ml, respectively) per 10 g body weight. Once under anesthesia, mice were fixed to the stereotaxic unit with head restrained using ear-bars. The head was shaved and an incision was made to expose the skull. Ten jeweler’s screws were inserted into the skull to support the microdrive implant. Two of these screws were soldered to wires and screwed until they pierced the Dura, which served as a ground/reference for EEG recordings (one in olfactory-bulb and one in the cerebellum). A 2 mm hole was made in the skull at position 1.8 mm medio-lateral and 1.8 mm antero-posterior to bregma junction. Tetrodes were lowered to about 0.7 mm from the surface of the brain (dura). Dental cement was spread across the exposed skull and secured to the microdrive. Any loose skin was sutured back in place to cover the wound. Mice were given Carprofen (5 mg/kg) post-operatively to reduce post-procedure pain. Mice usually recovered within a day after the tetrodes were lowered.

### Histology

To determine the exact final position of the tetrodes in a given animal we performed Nissl dye staining on hippocampal slices. After the experiment was concluded, mice were anesthetized with an overdose of 0.5 ml Ketamine and Xylazine solution (100 mg/ml and 15 mg/ml, respectively) and intra-cardiac perfused with 40 ml of 4% PFA solution, immediately after the tetrodes were retracted before mice were decapitated. The skull was cut open from the bottom of the head to expose the brain, which was gently removed and stored in 4% PFA solution for 24 hr in a shaker. The brain was coronally sliced in 30 μm thick sections using a cryostat at −13°C. The sections were stained with fluorescent Nissl dye (Neurotrace) and mounted onto a slide. The brain sections were viewed under a high magnification microscope, and digital pictures of the slices were taken.

### Experimental methods

After animals recovered their normal weight a few days post-surgery, the electrodes were connected to an amplifier to detect single cells. Tetrodes were lowered ~12.5microns twice a day (6 hrs. apart) until the number of recorded cells was maximized (this process took between seven to ten days). After the depth of the electrode was optimal, we waited 3-8 days for recording stabilization. Animals were food deprived for four hours before the recording sessions throughout the length of the experiment (7-35 days). Animal’s were reconnected to 16 or 32 channels for every single session of ten minutes and introduced in the same region of the arena relative to the recording room in each of the three environments. Each environment is approximately 50cm wide with walls that are 40 cm in height. All three environments present a cue card on the internal side of the enclosure with the same orientation relative to the room for all three environments (north on our plots in fig. 1) to act as a reference point for navigation. Environment A was a square enclosure with black walls and white floor that has ground food (vanilla cookie) spread over the surface of the floor and also a small pellet ~10cm away from the S-E corner of the environment. Environment B was a circular enclosure with black walls and brown textured floor with ground food (cocoa flavored puffed rice) spread over the surface. Environment C was a circular enclosure with white walls and floor and no food. We used these differences in shape and reward distribution to enhance the animal’s ability to differentiate between the environments. Animals were release to freely explore the environments and eat the food. For this paper, we only used the data from Environment A.

*Recordings* were done with Neuralynx amplifiers: Lynx-8, with a total of 16 to 32 channels depending on the success of the electrode’s implantation surgery. Data was collected in continuous format at a 30 kHz sampling rate and then pre-processed using NDManager (Hazan et al 2006). High and low frequencies were separated and spike wave-forms were extracted for cell identification. Cell clusters were computed using the EM algorithm implemented in the KlustaKwik software (KD Harris et al 2000) over the concatenated data of the 12 sessions within a day for each animal. Human inspection over each cluster was done using Klusters software (Hazan et al 2006). Spike times and animal trajectories were transformed into Matlab objects and data analysis was performed using this platform.

Before analyzing the data, the length of the useful portion of the experiment over days for a given mouse was determine based on our *criteria of recording stability*. Because we wanted “in principle” to follow the same cells over days, we established the following criteria for determine stability: for each cell we chose the channel with the largest spike-waveform fluctuation over the 4 channels on a given tetrode. We then computed over that channel the mean wave-form in a vector of 40 discretized points (at 30 kHz sampling rate). We calculated the peak amplitude of the mean waveform over all cells and also its standard deviation. We then compared the distributions of peak wave-forms for each cell across days using a t-test for two distributions with different means and standard deviations; when these distributions became statistically different we stopped using the data from that day on, for that particular animal.

### Data analysis

After units were isolated, cells were considered for the analysis if their average firing rate across the 4 sessions within a day was higher than 0.1 Hz. This threshold left out approximately 2% of the cells. To avoid spikes that are not correlated with mouse position due to pre-play or re-play (e.g. nonspecific spatial firing when animals are not moving within the box; Nádasdy et al 1999, Diba & Buszaki 2007, Davidson et al 2009), we only kept frames in which the animal was moving faster than 4 cm/sec. This threshold was obtained by scoring movies and avoid frames where animals were not moving their bodies but just their heads. We called these bins: “effective time bins”. For each spatial and angular bin (x,y,H, 3-dim, 10 bins per dimension, 1000 total spatial bins), we utilized the effective time bins to count the spikes occurring in each of bin. We then computed the mean-rate for each bin as the spike counts for each of the samples from each visit of the animal to a given spatial bin. We then divided the mean spike count by the size of the time bin (1/30 s) defined by our maximum accuracy for predicting animal position and heading-direction. Heading-direction (H) is the angle in 2-D between the position of the animal at time *t*_*i*_ and time *t*_*i*−1_. We also monitored the head direction of the animal measured by the positions of two LEDs attached to the pre-amplifier connected to the mouse tetrode boards, but this variable was not as well defined as the heading-direction (H) due to the many degrees of freedom of the animal’s head while moving during the session. We use the same definition of heading-direction as Sarel et al 2017.

### Model fitting

Our proposed model describes the angular dependency of the firing rate for all heading-direction angles and spatial bins. We binned each spatial and angular coordinate into 10 bins to have sufficient statistics within each bin. We used averages of 72 samples per bin during 40 minute recording per day. Position and heading-direction was computed at the video camera frame rate of 30 Hz. Heading direction was computed from the change in the animal’s position between two video recording frames (previous and current frame).

Because the model fit heading-direction modulation, we divided the firing rate for all three dimensions by the spatial dependence only, assuming that the modulation of the firing rate by heading-direction is multiplicative. Thus, we considered the quantity

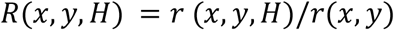

Over the entire article we only considered spatial bins in which average firing rates were larger than 0.5 Hz (we call this the “min-criteria”). We then used neurons that satisfy the following conditions: minimum number of spatial bins that satisfy the min-criteria = 10; minimum range of heading-direction angles to which the cell is responsive to = 50 deg. We also reproduced the same results with lower thresholds of minimum firing rate (0.1Hz).

The model has a periodic dependence on the angle between the heading-direction of the animal’s trajectory and the location of the reference point (See Fig.1b)

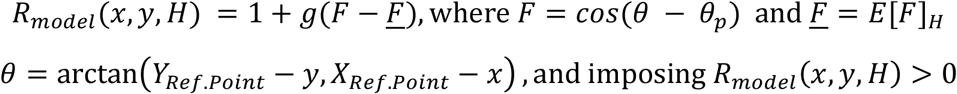

We fitted the values *g*, *θ*_*p*_, *X*_*Ref.point*_, *Y*_*Ref.point*_ by minimizing the variance of the model with respect to the data: *f* = [*R*_*model*_(*x,y,H*) − *R*(*x,y,H*)].

We minimized the variance utilizing an unconstrained nonlinear optimization procedure: the fminsearch function from MATLAB. For the initial parameter estimates we used: *g*=0;*θ*_*p*_=0; *X*_*Ref.point*_=Place-cell’s center of mass X-coord, *Y*_*Ref.point*_=Place-cell’s center of mass Y-coord.

### Statistical tests

In figure 1a, we computed the variance explained by a place description to describe how well the mean rate over all angular bins explain the neural activity,

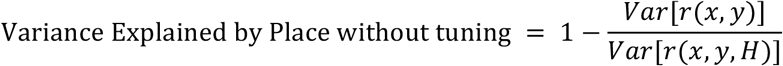

In figure 1b. we show, in a polar plot, the average rate for each of the 10 heading-direction angles, averaged over the entire experiment.

For figure 1d,e and f, we performed a chi-square goodness-of-fit test, and we rejected the hypothesis that the distributions in the X and Y axis were coming from the same distribution with the actual values computed being p_value_=6×10^−14^, p_value_=2×10^−36^, p_value_=0.028 respectively.

In figure 2b, we performed a chi-square goodness-of-fit test, and we rejected the hypothesis that the distributions in the X and Y axis were coming from the same distribution with the actual value computed being p_value_=4×10^−15^.

In figure 2c, we compared the variance explained by the place-tuning versus the variance explained by including reference-heading (RH) angular dependence where the reference point was fitted by the model:

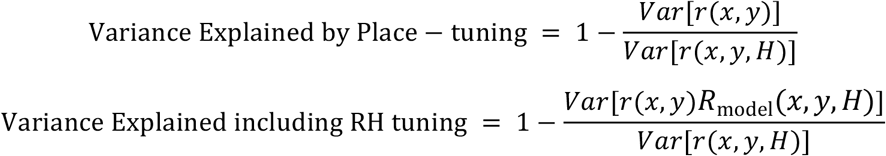

---

**Supplementary Figure 1:**
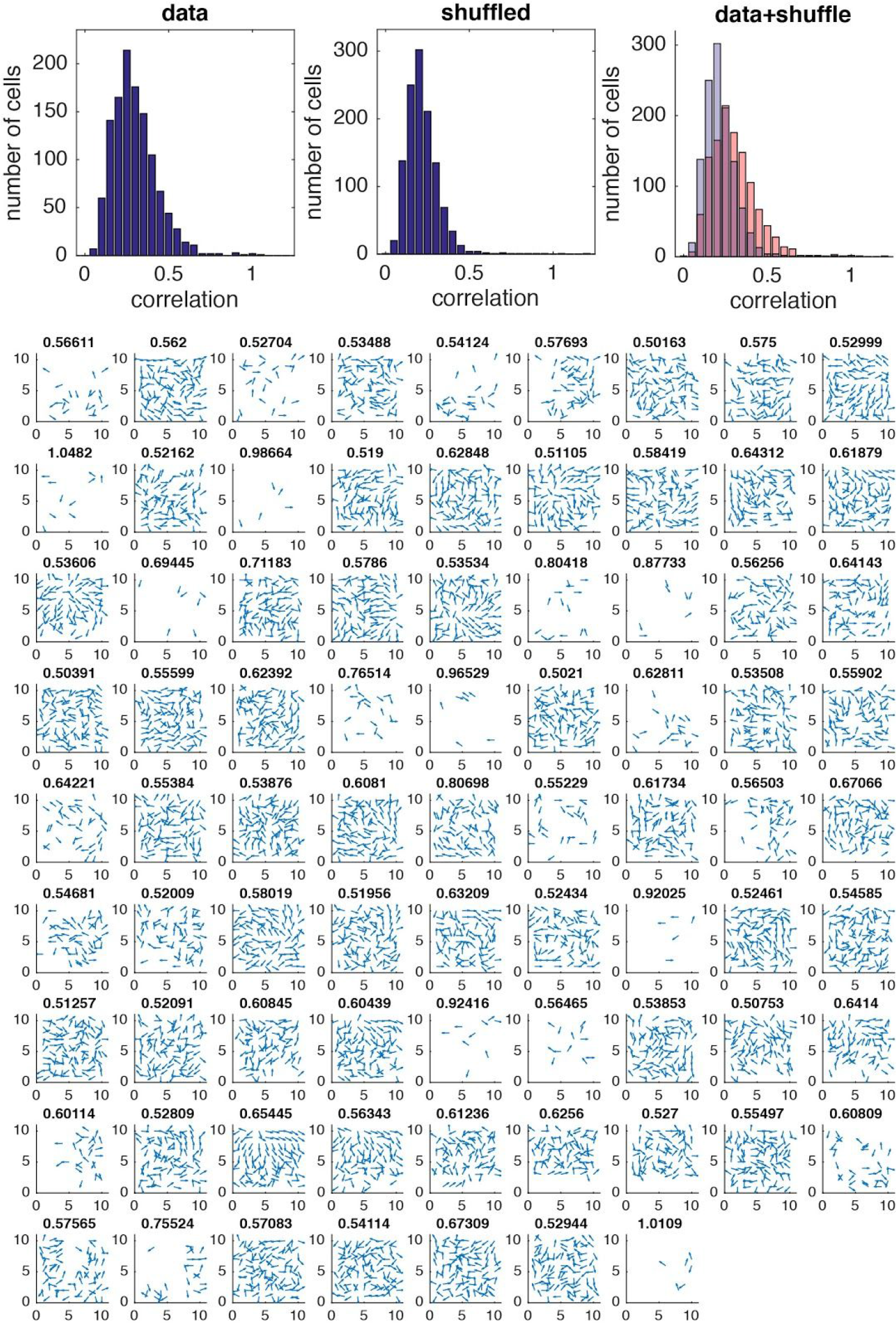
Normalized vector fields obtained from the heading direction with higher vector strength on each spatial bin. **Top**, Correlation of the vector fields for the raw data, and for shuffle data to compare the values of correlations that we would obtain at chance on a vector field of 10 by 10 bins. Correlation of the vector fields for the raw data versus shuffle, show a population of cells that could not be explain by randomly adding directions on the place cell field modulated by the behavior of the animal (pink region) **Bottom**, example of normalized vectors fields to visualize organization of prefer heading direction response. The structures in some neurons are global and give information of a perturbation like a curl, a divergence or a saddle-node at the other side of the arena.

**Supplementary Figure 2:**
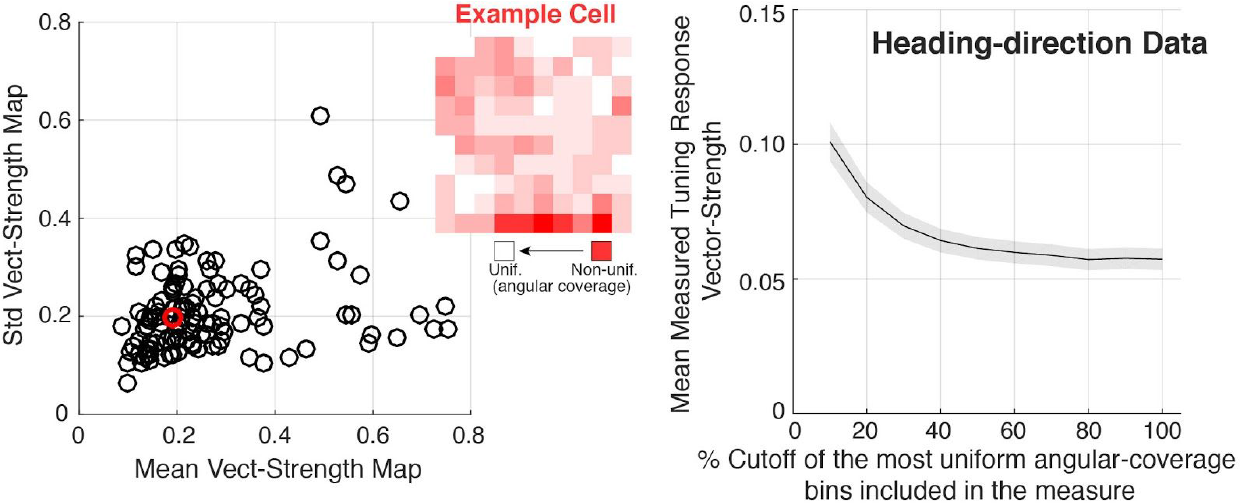
Non-uniformities in heading direction distribution per spatial bin, does not explain the angular tuning of neural responses. Each spatial bin has a distribution of heading directions manifesting the directionality of the trajectories when the animal visited that given spatial bin. **Left:** The angular distribution per bin is not uniform and is represented by the map on red tones on the top-left plot. The more red is the bin, the less uniform are the visits in a session of 40min. The distribution over the bins for the whole environment is represented by the mean and the std of distributions per bin, as a circle in the plot for all sessions. Red circle is a “average” sessions represented on the map above. **Right**: mean and +/− std of vector-strength for the populations of neurons when are included spatial bins with different levels of heading direction coverage, starting by the bins with the best 10% uniformity, until reaching the totality of bins indicated by 100%. As is it shown, the angular tuning decreases when bins with lower uniformity are included in the computation of the vector-strength. This rules out the argument by Mueller et al 1994, that the angular tuning in dorsal hippocampal neuronal responses is an artifact of the low angular coverage of certain bins of the environment close to the borders of the box. If any, including this bins into the computation of the angular tuning, decreases then tuning as it is shown here.

**Supplementary Figure 3:**
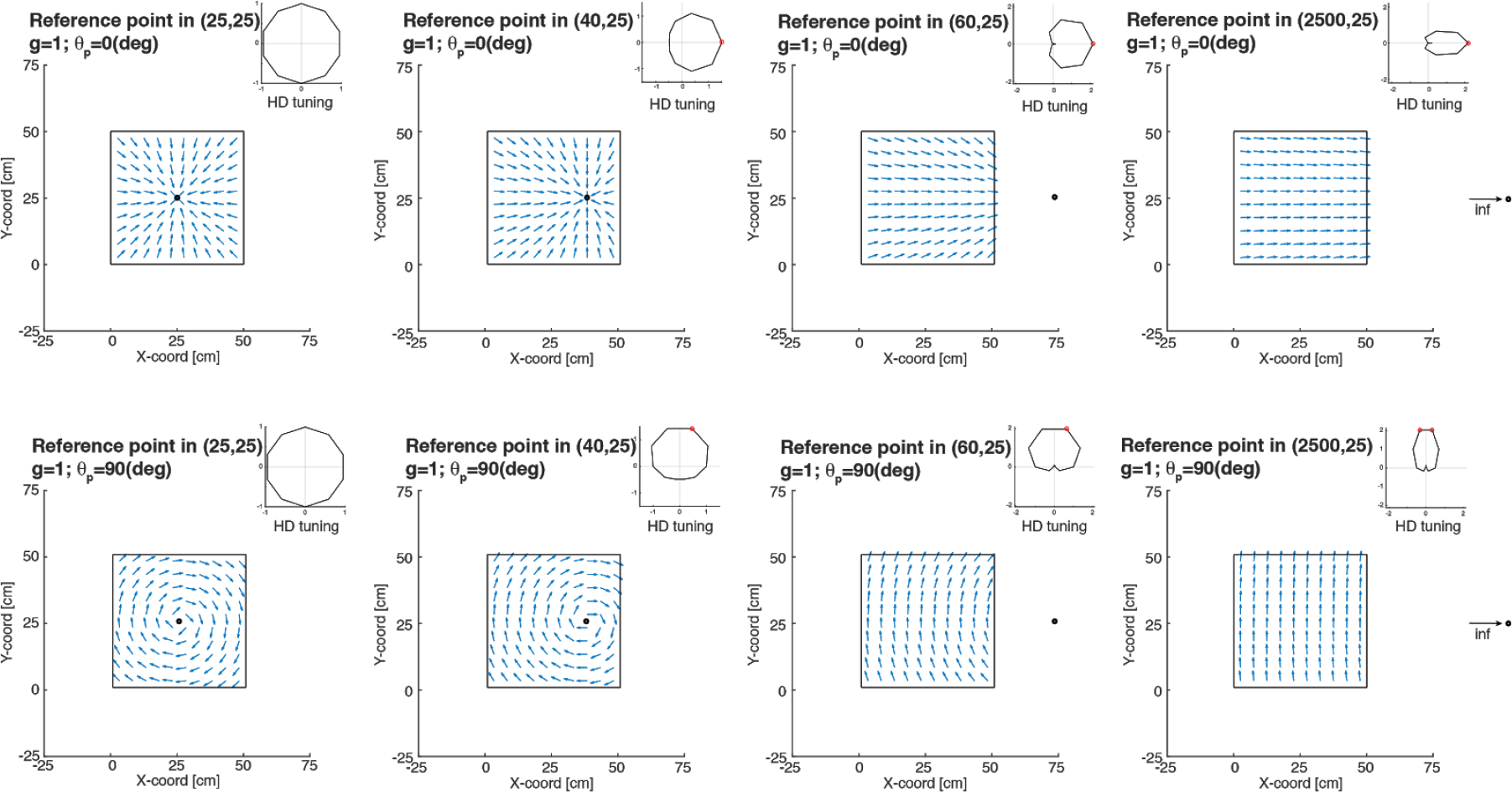
Examples of neuronal response for reference point tending to infinity generated by our model. **(Top-left)** If the reference point is within the arena and with phase angle (*θ*_*p*_ =0) neural response relative to the heading-direction is maximal when animal faces the reference point. **(Top-from-left to-right)** If the reference point moves from the center of the arena, the heading-direction vector field tend to point to the reference point reaching the point of a parallel field with reference point at the infinity. Heading-direction circular histogram shows the evolution towards a traditional pure Head-direction neuronal response. **(Bottom-from-left to-right)** as on the top row heading-direction response tend to behave as traditional head-direction cells when reference point moves to infinity. In these examples the *θ*_*p*_ is equal to 90, so the maximal neural response is when the animal moves perpendicular to the reference point.

**Supplementary Figure 4:**
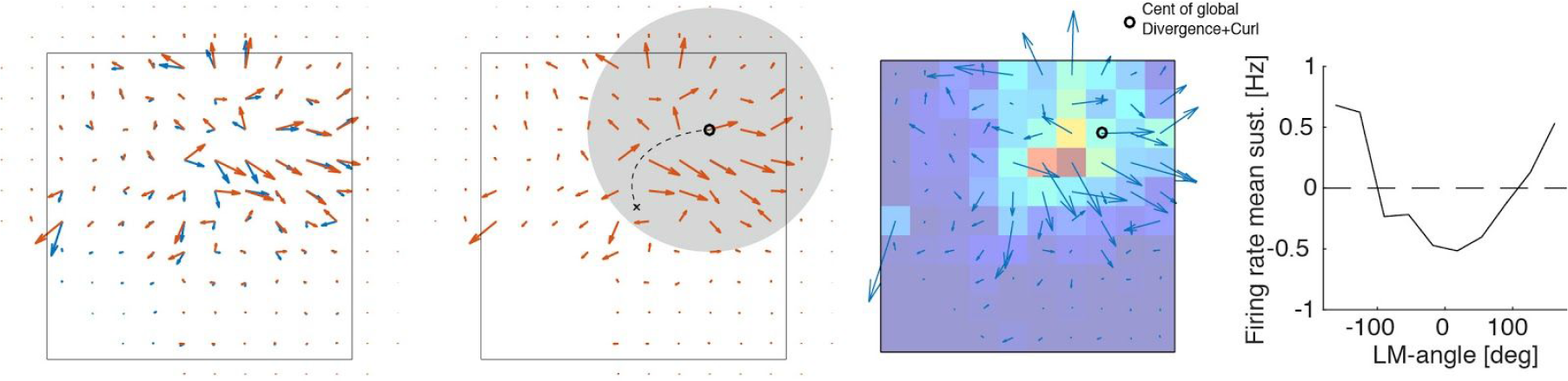
Computing the center of the divergence and the curl of the heading-direction responses’ vector field. **Far-left:** vector field generated by the vectors of the best heading direction responses in each bin (blue arrows) for all bins in the arena (10 by 10). Red arrows are the smoother version of the original vector field (blue) convolved with a 2-D gaussian filter for vector’s angle and length. This filter gives a 2 extra bins of edging. **Left:** once the vector field is smoothed, the location of the curl and the divergence (which are local descriptors of the field) are computed within a disc around the possible candidate, starting from the center of the arena and running in an iterative manner, in order to find the center of divergence+curl for a large part of the field. We will call this, the global center of divergence and curl for the neural response (black circle). **Right:** superposition of the heading direction vector field with the place-field in x-y coordinates. Far-right: neural response for the angle dependence relative to the center of the global divergence+curl.

**Supplementary Figure 5:**
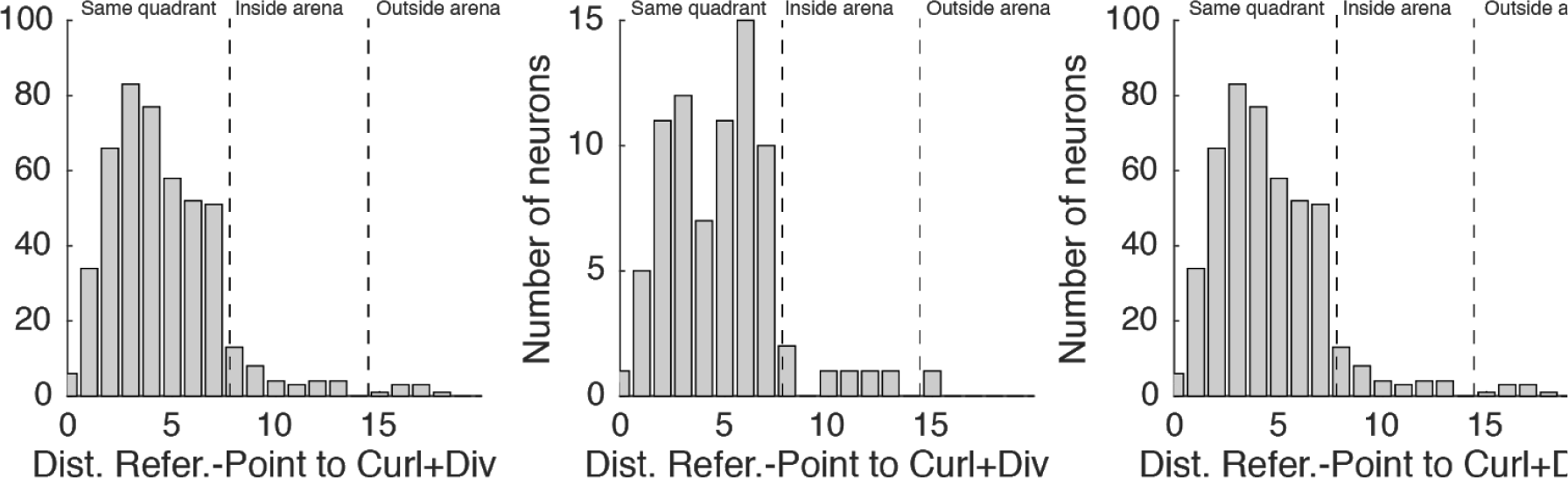
Curl+Divergence iterative method coincide with the reference point location obtained by our proposed model. The method described on Suppl.Fig.4 to find the Curl and Divergence of the heading-direction neural response provides candidates for the singularities related the reference points. The estimation of the reference point location from the two methods predicts this singular points to be within the same environment quadrant. The error between the methods is not small, but is mainly originated by the fact that we can only compute Curl and Divergence within the enclosure. We also show that this prediction is not random, since the distribution of the Curl+divergence distance to the place-fields center of mass (another comparable point within the enclosure) is much higher.

**Supplementary Figure 6:**
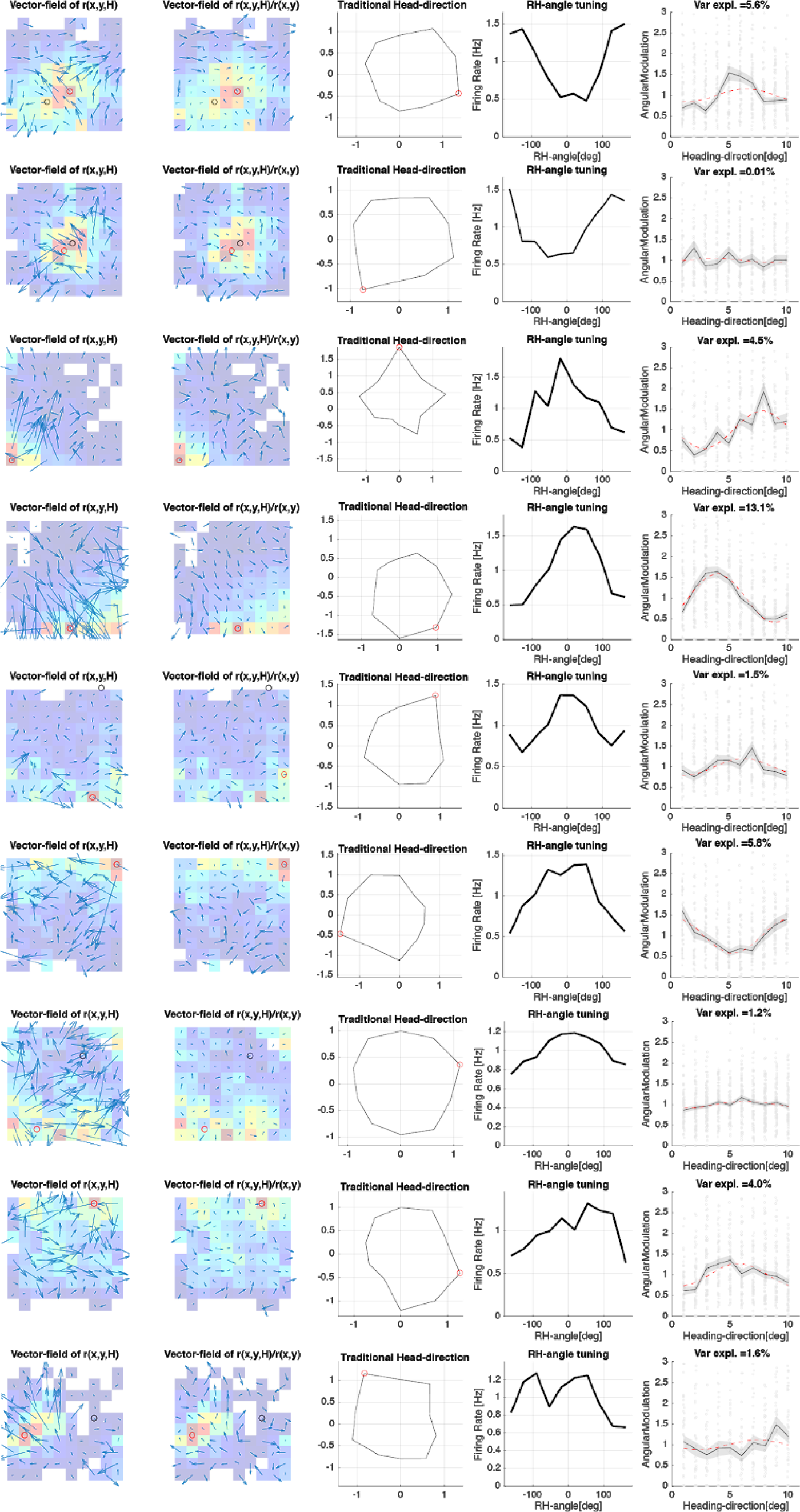
RH tuning for all cells included in Fig1c to show diversity in the RH-tuning responses.

